# Perceptual versus motor awareness of explicit contributions to visuomotor adaptation

**DOI:** 10.64898/2026.08.24.745320

**Authors:** S. Heirani Moghaddam, A. Decarie, R. Chua, E.K Cressman

## Abstract

In the current experiment, we compared reported perceptual awareness of the visuomotor rotation to motor awareness of changes in reaches established using the process dissociation procedure and drawing task following visuomotor adaptation to a large (50°; R50 group) or a small (30°; R30 group) cursor rotation. Results revealed that perceptual and motor awareness did not differ in magnitude for the R50 group and were significantly correlated. In contrast, while the R30 group perceptually reported being aware of the visuomotor rotation, motor awareness was significantly less and responses were not significantly correlated across tasks. Overall, results suggest that perceptual and motor tasks assess different processes underlying visuomotor adaptation to a small cursor rotation, such that perceptual awareness of the visuomotor rotation is not reflected in participants’ reaching performance on tasks assessing motor awareness.

## Introduction

When performing a goal-directed reach, the central nervous system transforms incoming sensory information related to the position of the hand and target into appropriate motor commands (Wolpert & Kawato, 1998). Visual and proprioceptive (i.e., felt position) cues can both be used to establish hand position (Jeannerod, 1988). In the laboratory, a mismatch between visual and proprioceptive signals related to hand position is created when participants reach in a virtual reality environment while a cursor on the screen misrepresents their actual hand position in space. For example, the cursor’s trajectory may be rotated or translated relative to hand motion (Baraduc & Wolpert, 2002; Cressman & Henriques, 2009; Heirani Moghaddam et al., 2021; Krakauer et al., 2000).

Participants typically experience reaching errors when initially reaching in a virtual environment with rotated cursor feedback, such that the cursor does not go to the target as expected. For example, if the cursor feedback is rotated clockwise (CW) relative to hand motion, participants see the cursor moving to the right of the target. However, participants quickly adapt their reaches (e.g., within 10 – 20 trials), reaching to the left of the target so that the cursor once again successfully lands on the target (Bastian, 2008; Krakauer et al., 2000; Maksimovic & Cressman, 2018). These leftwards reaches continue even when visual feedback is removed and participants are instructed to aim to the target (i.e., aftereffects), and are proposed to reflect implicit adaptation, arising in the absence of conscious awareness (Baraduc & Wolpert, 2002; Cressman & Henriques, 2009; Gastrock et al., 2020; Hegele & Heuer, 2010).

Explicit processes have also been suggested to play a role in visuomotor adaptation (Benson et al., 2011; Heirani Moghaddam et al., 2021; Maresch et al., 2020; Mazzoni & Krakauer, 2006; Modchalingam et al., 2019; Werner et al., 2015, 2019). Early efforts to establish the role of explicit processes in visuomotor adaptation used questionnaires at the end of the experiment to probe participants’ perceptual awareness of the visuomotor rotation (Benson et al., 2011). More recently, motor tasks such as the process dissociation procedure (PDP) and drawing task (DT) have been used to establish motor awareness of changes in reaches following visuomotor adaptation (Heirani Moghaddam et al., 2021; Neville & Cressman, 2018; Werner et al., 2015). Within Werner and colleagues PDP (adopted from Jacoby, 1991), participants’ awareness of changes in their reaches is determined by establishing differences in reaching performance when participants are instructed to reach in the absence of visual feedback while (1) using anything that they learned during rotated reach training trials (inclusion trials) and (2) not using what they learned during visuomotor training and reaching as they did during baseline trials (exclusion trials) (Heirani Moghaddam et al., 2021; Maresch et al., 2020; Modchalingam et al., 2019; Neville & Cressman, 2018; Werner et al., 2015, 2019). As for the DT, participants are asked to draw the trajectory their hand took to get the cursor to the target during reach training trials with the visuomotor rotation (Heirani Moghaddam et al., 2021; Ong & Hodges, 2010; Wijeyaratnam et al., 2022). Thus, this task is similar to the inclusion trials of the PDP, but participants complete the task with vision of their hand. To date, this task has not been used as extensively as the PDP.

While both perceptual and motor tasks have been used to establish explicit contributions to visuomotor adaptation, the relationship between awareness of the visuomotor rotation as established by perceptual reports and awareness of changes in one’s reaches as established by motor tasks is unclear. Recently, we (Heirani Moghaddam et al., 2021), compared reaching performance on the PDP to one’s responses on the post-experiment questionnaire (Benson et al., 2011) and reaching performance on the DT (Ong & Hodges, 2010; Wijeyaratnam et al., 2022) after participants adapted to a 40° visuomotor rotation. We found that the magnitude of participants’ motor awareness established via the PDP was correlated to their perceptual awareness of the visuomotor rotation and motor awareness of changes in their reaches as established by the PEQ and DT, respectively. Thus, results from Heirani Moghaddam et al. (2021) suggest that perceptual awareness of the visuomotor rotation and motor awareness of changes in reaches are related after adapting to a large cursor rotation.

In Heirani Moghaddam et al. (2021), perceptual and motor awareness were only compared after participants adapted their reaches to a 40° visuomotor cursor rotation. Explicit processes have been shown to contribute significantly to visuomotor adaptation for cursor rotations of 40° or more, regardless of method of assessment (Modchalingam et al., 2019; Neville & Cressman, 2018; Werner et al., 2015). In contrast, previous literature using the PDP method has shown limited motor awareness when the cursor rotation is 30° or less (Modchalingam et al., 2019; Neville & Cressman, 2018; Werner et al., 2015). It remains to be determined if this lack of awareness is consistent across perceptual and motor assessments following visuomotor adaptation to a small cursor rotation.

In the current experiment, we looked to establish the relationship between perceptual and motor awareness following visuomotor adaptation to both large and small cursor rotations. Results from this study will provide insight into if perceptual awareness of the visuomotor rotation is related to motor awareness of changes in reaches. Participants adapted their reaches to a large (50°; R50 group) or a small (30°; R30 group) visuomotor rotation and awareness was assessed using perceptual (PEQ) and motor tasks (PDP and DT). We hypothesized that the magnitude of perceptual awareness and motor awareness would be similar for the R50 group, but not for the R30 group. These results would suggest that conclusions drawn regarding the contribution of explicit processes in visuomotor adaptation to a small cursor rotation are dependent on method of assessment and that perceptual awareness of the visuomotor rotation is not reflected in awareness of changes in reaches.

## Methods

The procedures and a subset of the data presented below are part of a larger study examining visuomotor adaptation. Additional details regarding data related to the R30 group can be found in Decarie & Cressman, 2022.

### Participants

48 participants aged 18-38 years old (M = 23.8 years, SD = 1.3) were recruited for this study. Participants were right-hand dominant based on their reports on the modified version of the Edinburgh Handedness Questionnaire (Oldfield, 1971). Participants were randomly divided into two groups, a R50 group that reached with a large (50°) visuomotor rotation and a R30 group that reached with a small (30°) visuomotor rotation. All participants reported having no history of sensory, motor or neurological impairment, and all had normal or corrected-to-normal vision. All participants were naïve to the purpose of the study and had never participated in a visuomotor adaptation study involving reaching with rotated visual feedback in a virtual environment. Participant recruitment and data collection commenced after the Faculty of Health Sciences at the University of Ottawa approved a (COVID) Safe Research plan and ethical approval was attained from the University of Ottawa’s Health Sciences and Science Research Ethics Board. Prior to starting the experiment, all participants provided written informed consent.

### Apparatus

Testing took place using the KINARM End-Point Lab (KINARM Technologies, Kingston, ON) in a dark and secluded room. Participants were seated in a height-adjustable chair located in front of the experimental apparatus and grasped the handle of the KINARM using their right hand (Figure 1). The chair was positioned so that the participant’s forehead rested comfortably on the testing apparatus and they could reach to all the targets within the workspace to complete the task. The position of the chair was locked in place and maintained throughout the experimental session. Visual targets were projected from a downward facing monitor (LG 47LD452B-UA EzSign – 47” LCD TV; refresh rate: 60Hz, 2.6A; Seoul, South Korea) located 20.5 cm above a reflective surface that was located 20.5 cm above the robot handle (Figure 1A).

**Figure 1.**
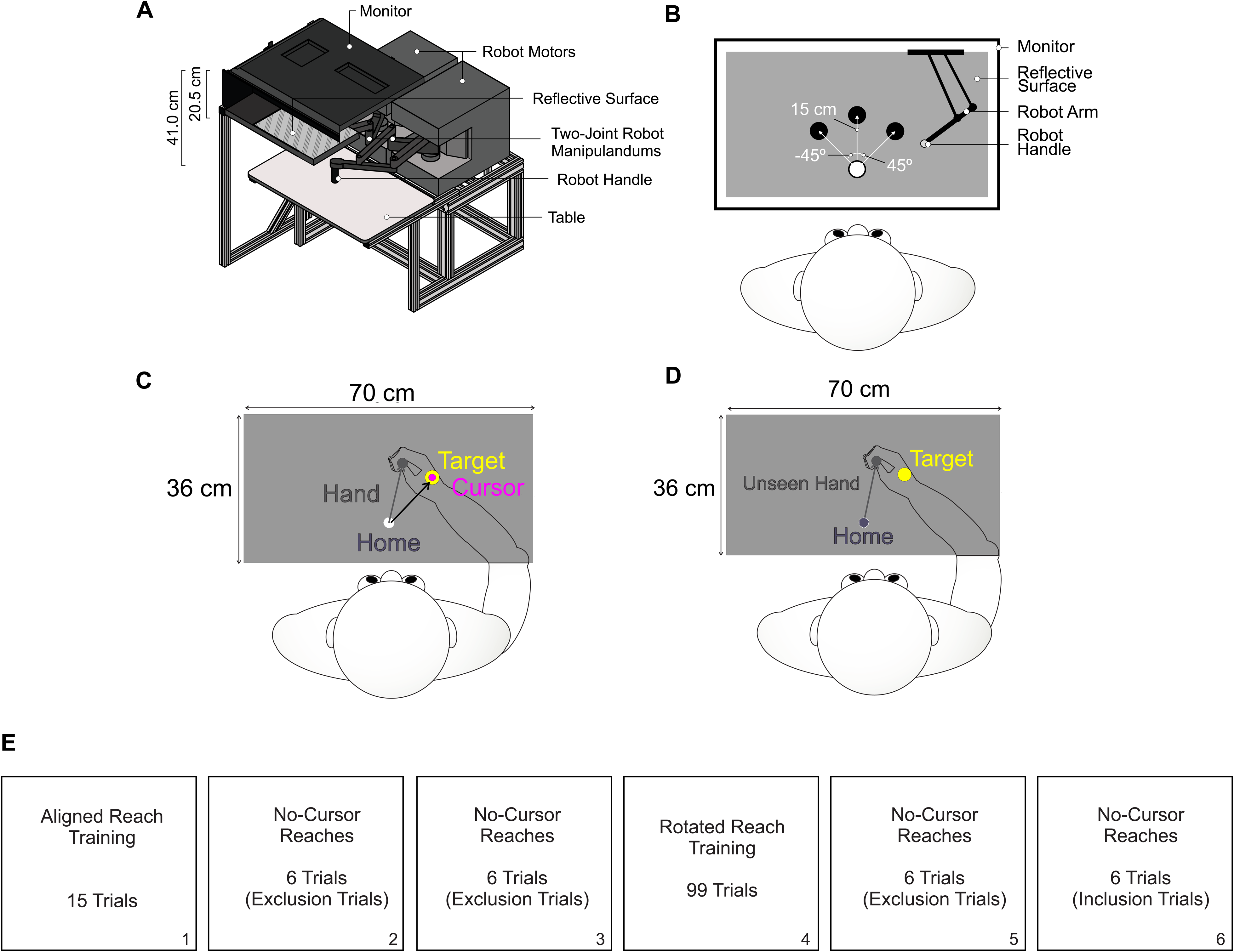
Experimental design including the apparatus and breakdown of testing blocks. **a.** Side-view of the experimental apparatus. Participants were seated in front of the apparatus and instructed to grasp the robot handle with their right hand. **b**. Top-down view of the 3 targets participants reached to during all reaching trials. **c.** Rotated Reach Training trials and **d.** No Cursor Reaches. **e.** Breakdown of testing blocks. The same blocks of trials were completed by both the R50 and R30 groups.

Thus, visual stimuli appeared to lie in the same horizontal plane as the right hand holding the robot handle. Participants’ view of their limbs was occluded by the reflective surface and a black cloth draped around their neck and attached to the apparatus. Once participants were seated comfortably, the room lights were turned off and testing began.

### Experiment Overview

Testing included a single session, which lasted approximately 1 hour. All participants completed the same blocks of reaches as outlined below (see Figure 1E).

For all reaches, the home (i.e., start) position (white circle, 2 cm in diameter) was located approximately 20 cm in front of a participant’s chest and in line with their body midline (Figure 1B). All movements began with the hand at the home position. Both groups first reached while seeing a cursor that was aligned with their hand (i.e., aligned reach training) to establish movement errors in a typical visual environment (baseline performance). Following aligned reach training, all groups then completed a block of no-cursor reaches to determine how they reached to a target when no visual feedback was provided (Figure 1D). Aligned reaches and no-cursor reaches were followed by participants reaching with rotated cursor feedback, where the cursor was rotated either 50° or 30° clockwise relative to hand motion depending on group (i.e., rotated reach training; Figure 1C). To assess the motor awareness of changes in reaches, a second block of no-cursor reaches were performed using the process dissociation procedure (PDP; described in detail below).

### Reach Training Trials

In aligned reach training (Figure 1E, Box 1), groups began by holding the robot handle at the home position for 500 ms. Following this, one of three targets appeared (yellow circle, 2 cm in diameter), located 15 cm away from the home position. The targets were located straight ahead of the home position (central target; 0°) and 45° to the left and right of the central target. Once a target appeared, participants were instructed to reach to it as quickly as possible with the goal of having the cursor land on the target. Real-time visual feedback of the hand position was provided via a cursor on the screen (magenta circle, 1 cm in diameter), both while the hand was held in the home position prior to the start of the reach, and throughout the duration of the movement. Once participants landed on the target (i.e., the center of the cursor and the center of the target were within 0.5 cm), the hand was held at this position for another 500 ms. The cursor and the target then disappeared, and the robot passively moved the hand back to the home position along a linear path in a movement time of 1000 ms. If participants attempted to move outside of the linear path, a resistance force (proportional to the depth of penetration with a stiffness of 2 N/mm and a viscous dampening of 5 N/mm) perpendicular to the grooved path was produced (Cressman & Henriques, 2009; Henriques & Soechting, 2003). Participants began the experiment by completing 15 aligned reach training trials (5 to each target). The position of the KINARM robot was recorded at a sampling rate of 1000 Hz, with a spatial accuracy of 0.1 mm. See Figure 2A for a timeline of events for aligned reach training.

**Figure 2.**
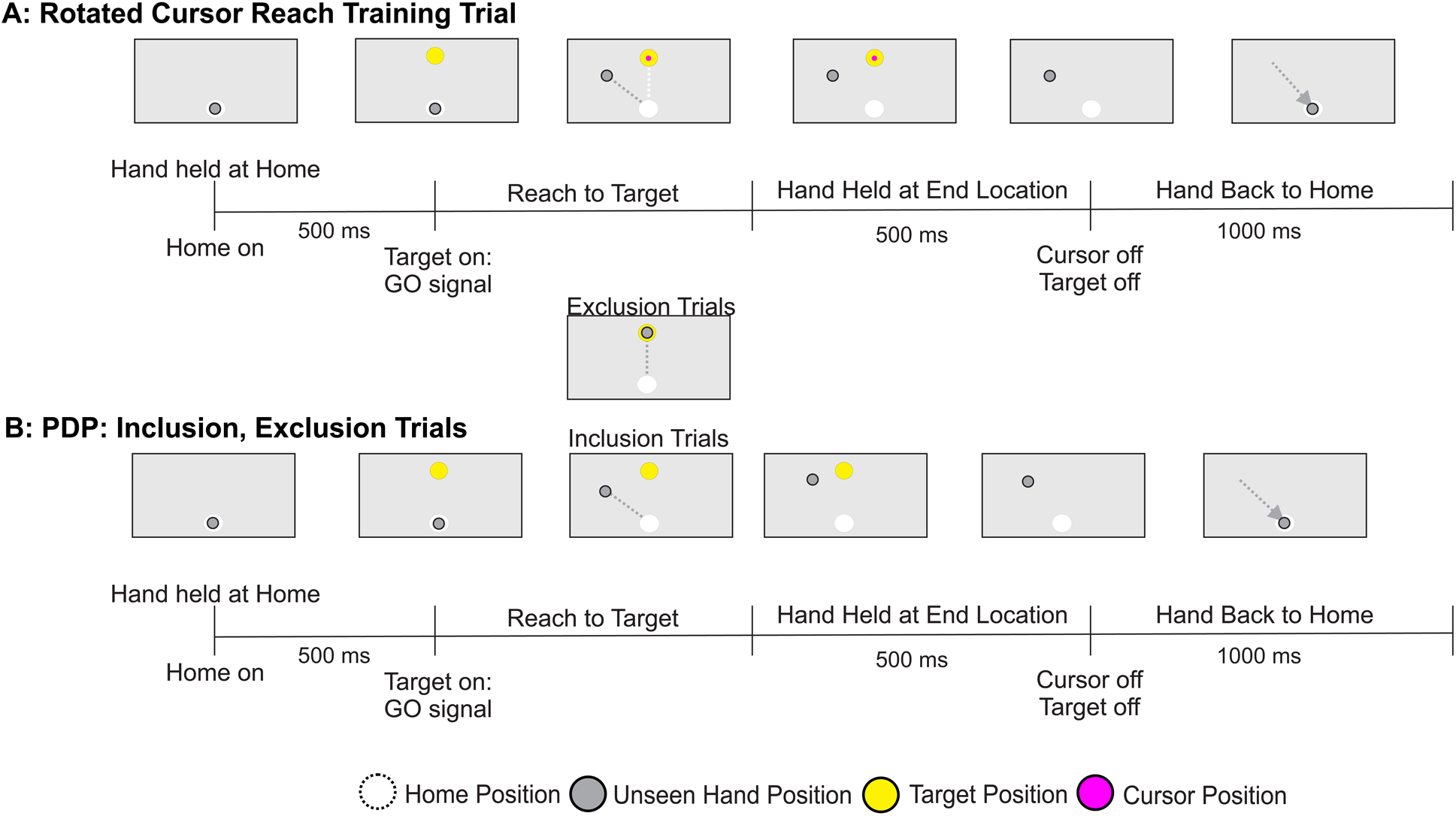
Timeline of events for each trial type. **a.** Rotated Reach Training **b.** PDP No-Cursor Reaches. In inclusion trials, participants were instructed to: “Reach using anything you have learned during training in order to get the cursor to the target. In other words, reach so that the cursor would have gone straight to the target, as in the reaching trials you just completed when the cursor was available”. In exclusion trials, participants were instructed to “Reach so that your hand goes straight to the target”.

The rotated reach training trials followed the same timeline of events as the aligned reach training trials explained above (Figure 2A), in that participants were instructed to reach to the target as quickly as possible with the goal of having the cursor land on the target. However, on these trials, the cursor representing the position of the hand was rotated 50° CW or 30° CW relative to the participant’s hand trajectory depending on which group they were in. Participants were not made aware of this rotation, nor were they given instruction on how to counteract it. Both groups performed 99 reaches (33 to each target; Figure 1E, Box 4) with the rotated cursor.

### No-Cursor Reaches to Assess Motor Awareness of Changes in Reaches

Two blocks of 6 no-cursor reaches (2 reaches to each target within each block) were performed twice: before (i.e., Time 1; Figure 1E, Box 2 and Box 3) and after rotated reach training (i.e., Time 2; Figure 1E, Box 5 and Box 6). For these trials, participants reached when no cursor was displayed (Figure 1D). These reaches followed the same timeline of events as described above in the reach training trials (see Figure 2C). However, the end of each movement was determined online as the time at which velocity first decreased below 0.01m/s. Using the PDP method, the no-cursor reaches at Time 1 included two blocks of *exclusion* trials, where participants were instructed to: “Reach so that your hand goes straight to the target”. Following the rotated reach training trials (Time 2), participants completed 6 *exclusion* trials. These were then followed by 6 *inclusion* trials, in which participants were instructed to: “Reach using anything you have learned during training in order to get the cursor to the target. In other words, reach so that the cursor would have gone straight to the target, as in the reaching trials you just completed when the cursor was available”. Participants always completed reaches under exclusion trials first, before they were cued about the presence of the visuomotor rotation in the inclusion trials, as it has been shown that implicit adaptation decays quickly (Bouchard & Cressman, 2021; Heirani Moghaddam et al., 2021; Neville & Cressman, 2018).

### Post-Experiment Questionnaire (PEQ) and Drawing Task (DT)

Once all reaching trials were completed, participants exited the testing room. They then completed the Post-Experiment Questionnaire (PEQ) of Benson and colleague (2011) designed to assess their ability to verbally report their perceptual awareness of the visuomotor rotation at the end of the experiment. Within the PEQ, participants were first asked whether they noticed a change during the experiment. Following a ‘yes’ response, they were then probed on how they would describe that change and asked to verbally provide an estimate of the size of the visuomotor rotation (i.e., indicate by how many degrees they had to move to the left of the target). These participants were designated as aware of the visuomotor rotation. If participants answered the first question by indicating ‘no’ they did not notice the task change, they were further asked if they noticed that the cursor did not move where they intended for it to move and if they had to correct their movements because of the cursor movement. If participants answered no to all questions they were designated as unaware of the visuomotor rotation (see Benson et al., 2011).

Finally, participants completed a drawing task (DT) in which the home position and three visual targets were drawn to scale on a sheet of paper. DT instructions consisted of 2 parts: 1) Think back to the final block of reaching trials you completed when the cursor was present on the screen and indicate **where your hand was** at the end of this movement, 2) Draw the **path your hand took** to get the cursor to the target (DT). A participant’s hand was visible during these trials. The order in which participants completed the PEQ and DT was counterbalanced across participants such that following the completion of all reaching trials, half of the participants completed the DT first followed by the PEQ, and the other half of the participants completed the tasks in the reverse order.

## Data Analysis

All reaching trials (i.e., aligned reach training trials, rotated reach training trials, and no-cursor reaches) were visually inspected using custom written programs for MATLAB. The start (i.e., movement onset) and end (i.e., movement termination) points of each movement were selected using a velocity-based criterion. Movement onset and movement termination were defined as when velocity first increased above, and decreased below, 0.01 m/s and remained above or below for 50 ms, respectively. For each trial, we determined the angular error of the hand relative to the target at peak velocity (i.e., PVAE), where PVAE is equal to the angular difference between a vector from the home position to the desired target and a vector from the home position to the hand’s actual position at peak velocity.

### Reach Training Trials

The extent of final reach adaptation achieved during rotated reach training was determined to be the change in average PVAE over the last 15 rotated reach training trials relative to the average PVAE of all 15 aligned reach training trials.

### Assessments of Awareness

#### Process Dissociation Procedure (PDP)

Explicit motor awareness of changes in reaches were established using the no-cursor reach trials. To establish motor awareness, we calculated the average PVAE of the 6 inclusion no-cursor reaches after rotated reach training (i.e., Figure 1E, Box 6) and subtracted the average

PVAE of the 6 exclusion no-cursor reaches after rotated reach training (i.e., Figure 1E, Box 5), according to the following formula:

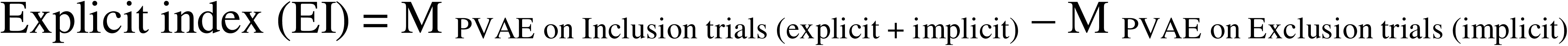

The average PVAE of the second block of 6 exclusion trials at Time 1 was then subtracted from EI values to account for baseline differences.

#### Post-Experiment Questionnaire (PEQ)

To analyze perceptual awareness of the visuomotor rotation as established via the PEQ, the angle reported by participants on the PEQ regarding changes in their hand trajectory was recorded. If participants reported that they did not notice any changes during rotated reach training trials, a score of zero was recorded.

#### Drawing Task (DT)

For the DT a reference line was first drawn between the home position and target. Then, the point on the drawn trajectory at which the greatest angular distance between the reference line and drawn trajectory was determined. From this point, a vector was drawn to home position. And the angle between the two vectors recorded. Responses were averaged across the trajectories drawn towards each of the three targets completed for each participant to establish motor awareness of changes in reaches.

## Statistical Analysis

Participants’ responses varied greatly within the different tasks for both the R50 and R30 groups. For example, with respect to motor awareness as established via the PDP, we found a standard deviation of 24.6° for the R50 group and 12.2° for the R30 group (see Supplementary File). In attempt to decrease variability across participants and given our previous work in which we used a subset of participants (see Heirani Moghaddam, Chua and Cressman, 2020), we included the 12 participants from both the R50 and R30 groups who demonstrated the greatest magnitude of motor awareness in the PDP trials in our analyses discussed below. That said, of these participants, 1 participant in the R50 group and 6 participants in the R30 group did not show evidence of motor awareness using the PDP (i.e., reaching errors on inclusion and exclusion trials were similar). These participants were kept in the analysis to ensure adequate sample number. Data for all participants is available in the supplementary file.

We performed Shapiro-Wilk tests to establish if the distribution of the data for all tasks (PDP, PEQ and DT) deviates from a comparable normal distribution and found non-significance (p > 0.05) for the R30 and R50 groups, indicating that the distribution of the samples for each group was not significantly different from a normal distribution. To normalize reach adaptation to the rotation size experienced, the extent of final reach adaptation was divided by 50° for the R50 group and 30° for the R30 group. These percentage data were then arcsine transformed before submitting to analysis. To determine whether the R50 and R30 groups adapted their reaches to a similar extent, we compared the arcsine transformed final reach adaptation values between groups using an independent samples t-test. Non-transformed values are reported below. The magnitudes of awareness as established via the PDP, PEQ and DT were then compared within the R50 and R30 groups. Specifically, repeated measures analyses of variance (RM-ANOVA) were completed with the factor of assessment method (PDP, PEQ, DT) for both the R50 and R30 groups. An additional ANOVA including the factor of order (participants who did DT first vs. DT second) was also completed for both the R50 and R30 groups. Given that we found no order effect, only results from the initial ANOVAs with the factor of assessment method are reported below. Further, correlational analyses were performed within each group to examine the relationship between the magnitudes of awareness as established via the PDP, DT and PEQ within participants.

The significance value for all statistical tests was set at *p* < 0.05, and Bonferroni post hoc tests corrected for multiple comparisons were used to find the locus of significant effects. When appropriate, Greenhouse–Geisser correction was applied and corresponding *F* and *p* values are reported. As well, coefficients of determination (R2), Pearson’s R, and slopes for linear regressions are reported to describe the relationship between the variables of interest where applicable.

## Results

### Reach Training Trials

The extent of final reach adaptation for both the R50 and R30 groups were similar in extent, such that the R50 group adapted their reaches by 42.0° or 84% (SD = 9%, 95% CI [39.5°, 44.6°]) of the 50° visuomotor rotation on average and the R30 group adapted their reaches by 24.7° or 82% (SD = 6%, 95% CI [23.7°, 25.7°]) of the 30° rotation on average. Independent samples t-test revealed no significant group differences in the extent of final visuomotor adaptation when expressed as a percentage of rotation size (*t*(22) = 0.728, *p* = 0.474, Cohen’s d = 0.297).

### Explicit Contributions to Visuomotor Adaptation

The magnitudes of awareness as assessed via perceptual and motor tasks are displayed in Figure 3 for both the R50 and R30 groups. With respect to the R50 group, ANOVA did not reveal a significant effect of assessment method (*F*(2, 22) = 0.538, *p* = 0.589, η2 = 0.032), indicating that perceptual awareness established via the PEQ (M = 29.3°, SD = 11.9°, 95% CI [22.1°, 36.1°]) and motor awareness established via the PDP (M = 24.3°, SD = 14.7°, 95% CI [16.0°, 32.6°]) and the DT (M = 24.0°, SD = 15.6°, 95% CI [15.2°, 32.8°]) did not differ in magnitude. For the R30 group, ANOVA revealed a significant effect of assessment method (*F*(2, 22) = 9.737, *p* < 0.001, η2 = 0.371). Post hoc analyses revealed that perceptual awareness established via the PEQ was significantly greater (M = 19.8°, SD = 12.5°, 95% CI [12.7°, 26.8°]) than motor awareness established via the PDP (M = 4.3°, SD = 9.0°; *p* = 0.006, 95% CI [-0.8°, 9.4°]) and DT (M = 0.7°, SD = 11.9°, 95% CI [-6.1 °, 7.4°]; *p* < 0.001). Furthermore, motor awareness as assessed via the PDP and DT did not differ from each other (*p* = 1.00), nor 0 (PDP: *t*(11) = 0.9704, *p* = 0.35; DT: t(11) =-0.0434, *p* = 0.97).

**Figure 3.**
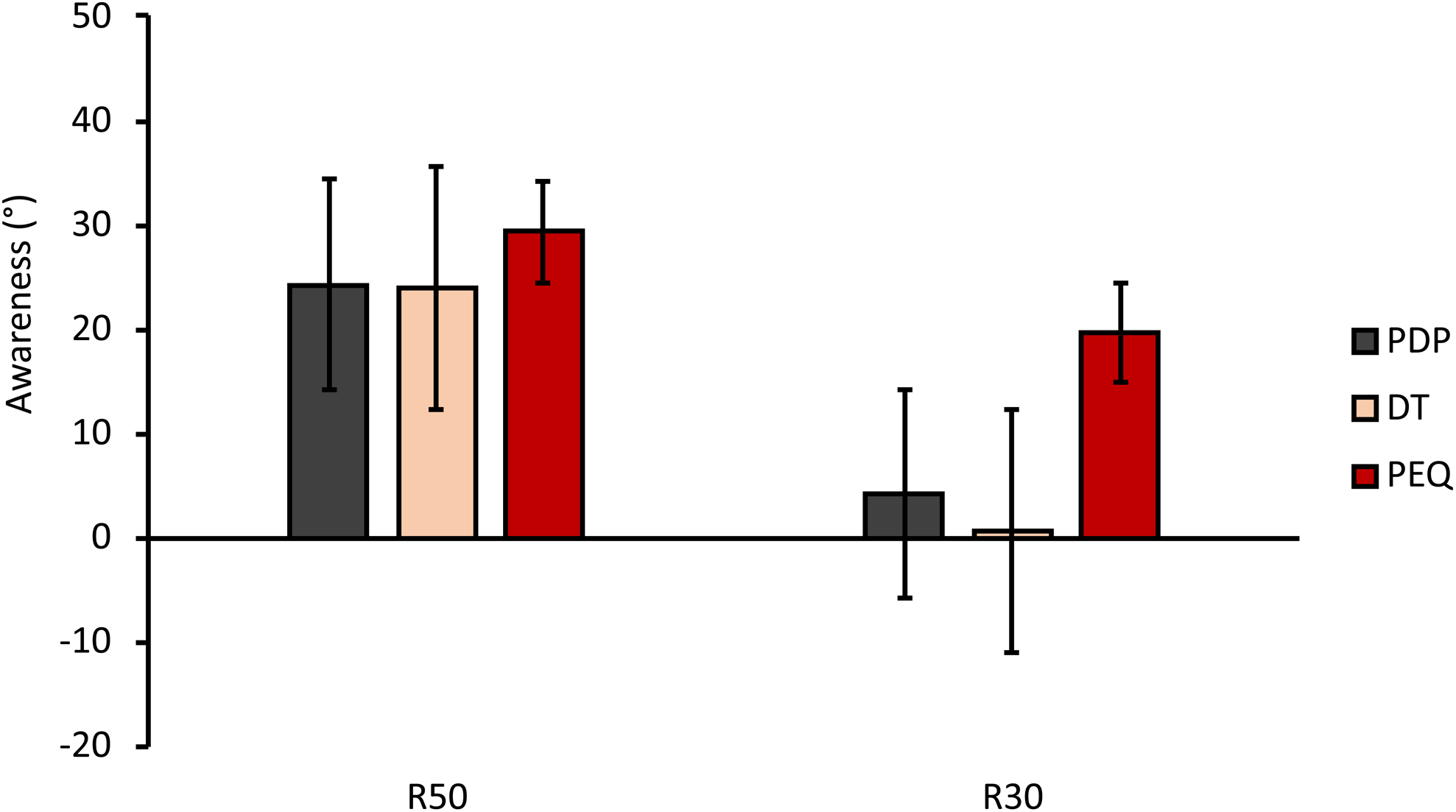
Awareness. Data for the R50 group is displayed on the left, and data for the R30 group is displayed on the right. All data presented reflect the magnitude of awareness as established via the PDP (black bars), DT (beige bars) and PEQ (red bars). Error bars represent the standard error of the mean.

The relationship between awareness established at an individual participant level across methods of assessment is shown in Figure 4 and Table 1. In the R50 group, Pearson correlation analyses revealed a significant correlation between motor awareness established via the PDP and DT (*p* = 0.005), PDP and PEQ (*p* = 0.005) but not the DT and PEQ (*p* = 0.132; see Table 1 and Figure 4 for details). In the R30 group, Pearson correlation analyses revealed a significant correlation between the two motor tasks used to assess motor awareness in the R30 group (*p* = 0.045; see Table 1 and Figure 4 for details). However, responses on the PEQ were not significantly correlated to either of the motor tasks for the R30 group (both *p*s > 0.05).

**Figure 4.**
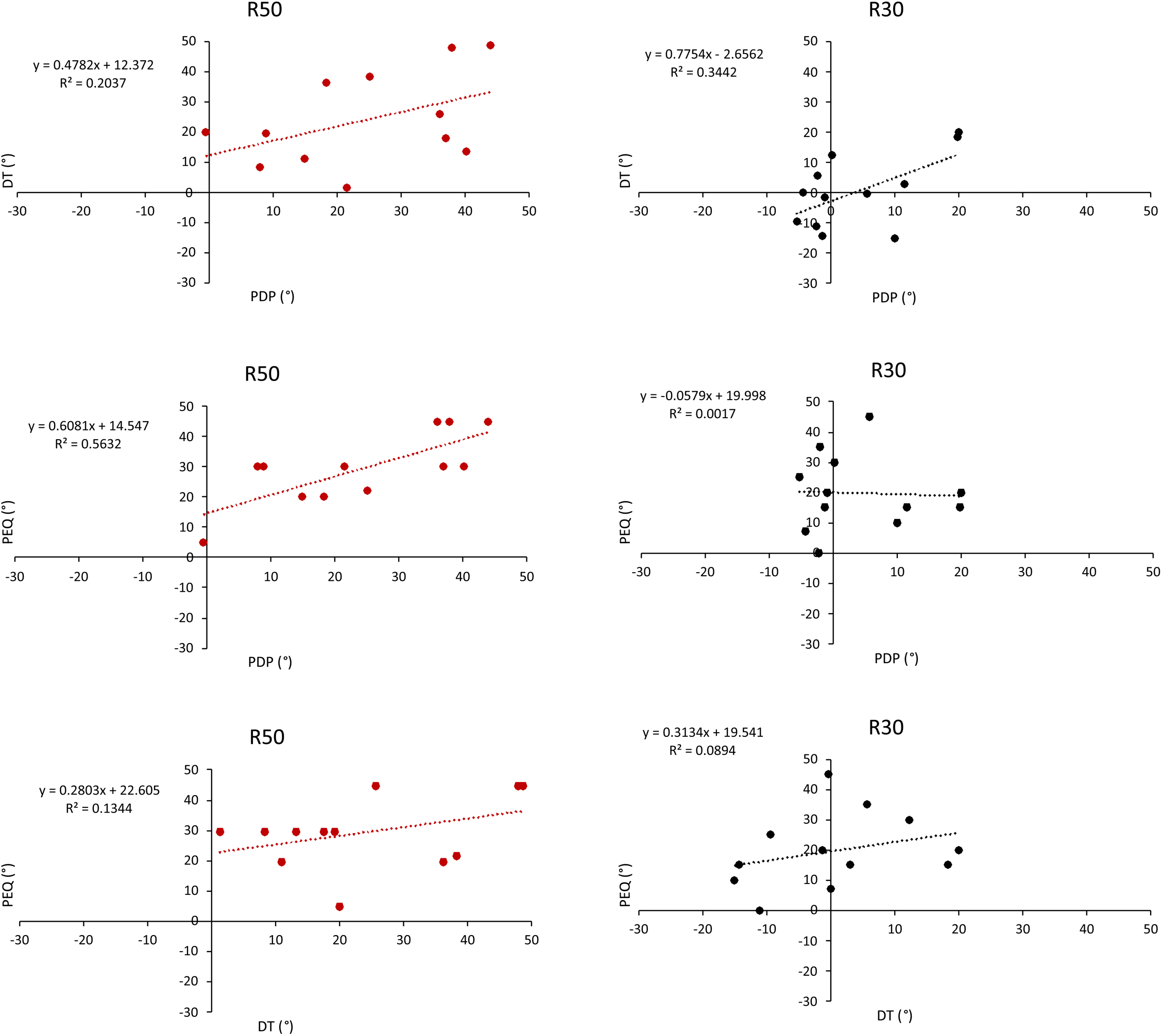
Relationship between awareness established via perceptual and motor tasks. Data for the R50 group are presented on the left (red dots) and data for the R30 group on the right (black dots). Respective equations for each line, R^2^ and *p* values are included on each plot.

**Table 1.** Pearson’s correlation results depicting the relationship between awareness as assessed via perceptual (PEQ) and motor tasks (PDP, DT) for the R50 (top) and R30 (bottom) groups.

## Discussion

In the current experiment, we compared perceptual awareness of the visuomotor rotation to motor awareness of changes in reaches following visuomotor adaptation to a large (50° (R50 group)) and a small (i.e., 30° (R30 group)) visuomotor rotation. Within group analyses revealed that for the R50 group, awareness established via perceptual and motor tasks did not differ in magnitude and motor awareness established via the process dissociation procedure (PDP) was significantly correlated with perceptual and motor awareness established via the post-experiment questionnaire (PEQ) and drawing task (DT), respectively. In contrast, while the R30 group reported perceptual awareness of the visuomotor rotation using the PEQ, the PDP and DT revealed a lack of motor awareness. Furthermore, correlation analyses revealed non-significant relationships between perceptual awareness established via the PEQ and both motor tasks for the R30 group. Overall, results suggest that perceptual and motor tasks assess different aspects of awareness following adaptation to a small visuomotor rotation, such that perceptual awareness reported via the PEQ is not reflected in motor awareness assessed via the PDP or DT.

### Method of assessment

Numerous assessments have been put forth to examine explicit contributions to visuomotor adaptation. These methods include having participants perform post-experiment questionnaires (e.g., PEQ; Benson et al., 2011), reach in the absence of visual feedback while engaging or not engaging in a strategy (e.g., PDP; Heirani Moghaddam et al., 2021; Modchalingam et al., 2019; Neville & Cressman, 2018; Werner et al., 2015, 2019), perform drawing tasks (DT; Ong & Hodges, 2010), report pre-planned aiming strategies prior to reaching to the target ((Bond & Taylor, 2015; Taylor et al., 2014)), and fixate at a pre-planned aiming point prior to reaching (de Brouwer et al., 2018). In the current experiment, we established perceptual awareness of the visuomotor rotation using the PEQ and motor awareness of changes in reaches using the PDP and DT after participants adapted their reaches to a 50° or 30° cursor rotation.

With respect to the R50 group (N = 12), average motor awareness was 24° as determined by the PDP and 92% of participants included in the analysis showed evidence of motor awareness. As well, 100% of participants in the R50 group were able to verbally report awareness of the visuomotor rotation using the PEQ and showed evidence of motor awareness within the DT. These results suggest that the large 50° cursor rotation was sufficient to elicit changes in reaches that could be accessed by the motor and perceptual systems following visuomotor adaptation. Overall, these results are consistent with our previous results demonstrating the extent of motor and perceptual awareness following visuomotor adaptation to a 40° cursor rotation was correlated across the PDP, DT and PEQ (Heirani Moghaddam et al., 2021).

In contrast to the R50 group, the R30 group exhibited very minimal motor awareness of changes in reaches (4°) when assessed via the PDP, despite clear evidence of visuomotor adaptation to the 30° cursor rotation. Looking at individual participant data in more detail, only 50% of the 12 participants included in our analysis for the R30 group showed evidence of motor awareness established via the PDP (and only 6 out of the total 24 participants; see Supplementary File for individual participant data). With respect to the DT, 42% of participants in the R30 group demonstrated evidence of motor awareness (and only 7 out of the total 24 participants) and reaching errors on the DT were correlated with reaching errors in the PDP. This limited engagement of strategic reaching and hence motor awareness observed when adapting to a small cursor rotation is in agreement with previous results of Heuer and Hegele (2014). In their paradigm, older adults demonstrated impaired visuomotor adaptation to large cursor rotations, when strategic corrections are typically engaged (Heuer & Hegele, 2014).

While limited motor awareness was established for the R30 group, 96% of participants in the R30 group reported perceptual awareness of the visuomotor rotation at the end of the experiment using the PEQ (19 out of the total 24 participants). Overall, results suggest a large variability between participants’ perceptual and motor awareness as established across methods of assessment when participants adapted to the small 30° distortion. Furthermore, perceptual awareness established via the PEQ following adaptation to a small visuomotor distortion was not related to motor awareness as established via the PDP and DT.

The notion that different methods of assessment are not measuring similar explicit processes underlying visuomotor adaptation has been suggested previously (Heirani Moghaddam et al., 2021; Maresch et al., 2020, 2021). For example, evidence suggests that perceptual reports related to a pre-planning aiming strategy (see Taylor et al., 2014), are not related to motor awareness established using a motor task such as the DT after adapting to a large cursor rotation (Heirani Moghaddam et al., 2021). Moreover, Maresch et al. (2020) found a difference between the magnitudes of awareness established via perceptual and motor methods of assessment, such that participants pre-planned aiming strategies reflected a perceptual awareness of 17.8° (i.e., 30% of the 60° visuomotor rotation introduced) compared to a motor awareness established via the PDP of 1.2° (i.e., 2% of the 60° visuomotor rotation introduced). Based on these results, Maresch and colleagues (2020) concluded that perceptual reports provide a higher estimate of awareness compared to motor-based assessments. This is similar to the results found in the current experiment where the R30 group showed a higher magnitude of perceptual awareness compared to motor awareness assessed via the PDP and DT. Further, other experiments have used the PDP method to examine motor awareness following visuomotor adaptation to a small visuomotor rotation and found very small magnitudes of motor awareness (Neville & Cressman, 2018; Werner et al., 2015). Specifically, Neville and Cressman (2018) found 0.274° of motor awareness (1.4% of the 60° visuomotor rotation introduced).

### Perception and action visual processing model

The dissociation between awareness as established via perceptual and motor responses, at least when adapting to small cursor rotations, may be explained by Milner and Goodale’s dual visual processing model for action and perception. According to the dual visual processing model, neural substrates underlying visual perception differ from those underlying the visual control of goal-direction actions (Jacoby, 1991; Reingold & Merikle, 1990; Shanks & St John, 1994). Specifically, the ventral visual stream transforms visual information into perceptual representations that encapsulate spatial relations between objects, while the dorsal visual stream mediates the visual control of skilled actions such as reaching (Milner & Goodale, 1993, 2008). In the current experiment, perceptual awareness of the visuomotor rotation was established via verbal responses (perceptual task) in the PEQ. In contrast, motor awareness was established based on reaching performance (action task) within the PDP and DT. Perhaps the PEQ engaged a perceptual visual representation, whereas the motor tasks engaged an action visual representation. For both the R50 and R30 groups, we found a significant correlation between explicit adaptation as established via the PDP and DT, supporting the notion that a similar action visual representation mediated responses across both of these motor tasks.

For the R30 group, results from the PEQ suggest that while participants were aware that something changed during the reach training trials, the motor system was not aware or did not have access to this information. Within Milner and Goodale’s dual visual processing model it is argued that the perceptual ventral visual stream represents our visual experience of the world, but it does not provide a direct foundation for action (Milner & Goodale, 2008). As well, while the ventral stream identifies and plans goal-directed reaches, the implementation of the action is driven by the dorsal visual stream (Milner & Goodale, 2008). Therefore, following visuomotor adaptation to a small cursor rotation, the motor system may not be able to access the perceptual knowledge or awareness that a rotation was presented during reach training. Thus, reaching errors are similar across inclusion and exclusion trials, as participants cannot turn their perceptual awareness on as needed within the PDP (and DT).

### Limitations of methods of assessment

The present results do not allow us to draw conclusions regarding the best approach to assess explicit contributions to visuomotor adaptation. Instead, the results allow us to investigate the relationship between perceptual and motor assessments of awareness. As indicated below, limitations are present across assessment methods and comparing between assessment methods.

Perceptual tasks that rely on verbal reports to establish awareness of the visuomotor rotation have been criticized for underestimating awareness, since participants’ responses may be held with low confidence, biased by feelings of familiarity or impacted by differences in retrieval context (Jacoby, 1991; Mandler, 1980; Maresch et al., 2020, 2021; Reingold & Merikle, 1990; Shanks & St John, 1994). Interestingly, we found the greatest magnitude of awareness in the R30 group when perceptual awareness was assessed with the PEQ. In contrast to our results, Modcahlingam et al. (2019) found minimal evidence of perceptual awareness using the PEQ after visuomotor adaptation to a 30° rotation. That said, their participants were not asked to perceptually report a number to index the rotation size experienced, rather participants were assigned awareness levels of “None”, “Low” or “High” and received corresponding scores of 0, 1 and 3, respectively (Modchalingam et al., 2019). In the current experiment in which participants had to report the amount of the perceived rotation experienced, we observed large perceptual awareness following visuomotor adaptation to a small 30° rotation. It is important to keep in mind that responses for our participants on the PEQ were still less than the rotation size experienced by both the R50 and R30 groups (approximately 59% and 66% of the 50° and 30° rotation experienced, respectively). Therefore, participants in the current experiment still reporting limited perceptual awareness at the end of the experiment using the PEQ.

In the current experiment we categorized the DT as providing an assessment of motor awareness. The DT has not been used as extensively as the PEQ or PDP (see Heirani Moghaddam et al., 2021; Maresch et al., 2021; Ong & Hodges, 2010), and one could question if it assesses motor or perceptual awareness. We categorized the DT as a motor task given that we were requiring participants to perform a goal directed movement to establish awareness of changes in their reaches. That said, participants had vision of their limb readily available throughout this assessment and it was completed at the end of the experiment, in a different environment, requiring participants to recall changes in their reaches. These features are typically associated with perceptual tasks (Glover & Dixon, 2002; Haffenden et al., 2001; Plodowski & Jackson, 2001). In support of our categorization of the DT as a motor task, results of our correlational analyses suggest that for both the R50 and R30 groups, awareness established via the DT was correlated with motor awareness established via the PDP.

Within the PEQ and DT, we asked participants to provide an estimate of angular errors related to the visuomotor distortion and their hand path to establish perceptual and motor awareness respectively. Interestingly, we found no correlation between estimates provided in the PEQ and reaching errors in the DT for either the R50 and R30 groups, indicating that the angular errors verbally reported where not reflected in their reaches. While preliminary work from our own lab has demonstrated that participants are fairly accurate at estimating angles between two visual lines (average absolute error = 4°; (Heirani Moghaddam & Cressman, 2021), future work is required to establish the accuracy with which participants can report a visuomotor distortion, and the relationship of these estimates across tasks (e.g., PEQ and DT).

In contrast to the PEQ and DT, the PDP was completed in the same testing environment as the reach training trials. Within the PDP participants were asked to reach while engaging (inclusion trial) or not engaging (exclusion trials) in what they had learned (PDP). Motor awareness of changes in reaches was determined by the difference in reaching errors between inclusion and exclusion trials. Thus, the PDP assumes that participants are able to turn a reaching strategy on and off and that implicit process(es) engaged are the same in both inclusion and exclusion trials. This assumption of consistent implicit engagement has been challenged by experiments suggesting that multiple implicit processes underlie visuomotor adaptation and may be differentially engaged across types of trials (Heuer & Hegele, 2015). Recent studies have also suggested multiple explicit processes underlie visuomotor adaptation (Maresch et al., 2020; McDougle & Taylor, 2019), including 1) a re-calculated component for every reach trial when reaching with a visuomotor distortion and 2) a memorized component that does not need to be re-calculated for every reach training trial. Future research is required in order to understand the relationship between multiple implicit and explicit processes across different trial types.

A final consideration across all methods of assessment is the variability of responses between participants. Specifically, standard deviation of perceptual awareness across participants as assessed via the PEQ was 11.9°, and motor awareness as assessed via PDP and DT were 14.7°, and 15.6°, respectively. Conversely, we found that in both R50 and R30 groups, standard deviation of the magnitude of implicit adaptation as established via the exclusion trials of the PDP was more consistent between participants. Specifically, in the R50 group the standard deviation of implicit adaptation was 5.5° with a 95% CI of [9.0°, 15.2°] and for the R30 group the standard deviation of implicit adaptation was 4.2° with a 95% CI of [9.5°, 14.2°]. Future work is required to establish sources of variability within a participant’s responses and across participants’ responses for the various awareness assessments (e.g., look at impact of instructions, time of evaluation, etc.).

Recognizing these limitations, the goal of this experiment was to investigate the relationship between methods of assessment that establish perceptual and motor awareness following visuomotor adaptation to a small and large rotation. Our results indicate that perceptual and motor awareness were similar in the R50 group whereas perceptual awareness as established via the PEQ was significantly larger than motor awareness in the R30 group.

## Summary

In the current experiment, we examined the relationship between awareness as assessed via perceptual and motor tasks following visuomotor adaptation to a large and small cursor rotation. We found that following adaptation to a large (i.e., 50°; R50 group) cursor rotation, perceptual awareness of the visuomotor rotation and motor adaptation of changes in reaches were similar in magnitude and correlated with each other. In contrast, following adaptation to a small (i.e., 30°; R30 group) cursor rotation, the magnitude of perceptual awareness established via the PEQ task was significantly larger than the magnitudes of motor awareness established via the PDP and DT. These findings support the idea that the PEQ is an unreliable measure of perceptual awareness since PEQ responses are not reflected in motor awareness, especially for smaller rotation sizes. Moving forward, future work needs to be careful with interpreting PEQ results and overall, consider the finding that different manners of assessment lead to variations in the extent of awareness established, specifically for a small cursor rotation.

