## Supplemental Table 1 for "Perceptual versus motor awareness of explicit contributions to visuomotor adaptation"

Authors: Heirani Moghaddam, S., Chua, R., Decarie, A., Vinh, M., Apreutesei, D., Cressman, E.K.

| Group: R50  Data provided for all 24 participants  Participants included in the analyses are indicated in grey (light grey across rows)  Positive values indicate reaching errors to the left of the target (in the expected direction of reach adaptation)  * outliers removed based on PEQ (pink) | | | | | |  |
| --- | --- | --- | --- | --- | --- | --- |
| Pnum | Final Adaptation | Final Adaptation ARCSINE Transform | Process Dissociation Procedure (PDP) | Drawing Task (DT) | Post Experiment Questionnaire (PEQ) | Implicit (PDP) |
| R50_21 | 83.7 | 66.2 | 44.0 | 48.7 | 45 | 6.4 |
| R50_12 | 70.1 | 56.8 | 40.2 | 13.3 | 30 | 16.1 |
| R50_17 | 52.9 | 46.7 | 38.0 | 48.0 | 45 | 1.4 |
| R50_7 | 82.4 | 65.2 | 36.9 | 17.7 | 30 | 10.1 |
| R50_13 | 67.4 | 55.2 | 36.1 | 25.7 | 45 | 9.8 |
| R50_14 | 87.3 | 69.2 | 25.2 | 38.3 | 22 | 12.6 |
| R50_8 | 75.1 | 60.1 | 21.5 | 1.3 | 30 | 7.6 |
| R50_15 | 57.1 | 49.1 | 18.3 | 36.3 | 20 | 11.4 |
| R50_24 | 62.6 | 52.3 | 15.0 | 11.0 | 20 | 18.0 |
| R50_4 | 79.5 | 63.1 | 9.0 | 19.3 | 30 | 18.5 |
| R50_5 | 94.7 | 76.7 | 8.0 | 8.3 | 30 | 20.0 |
| R50_20 | 93.9 | 75.7 | -0.5 | 20.0 | 5 | 13.0 |
| R50_19 | 50.7 | 45.4 | - 7.4 | -2.7 | 30 | 8.9 |
| R50_06 | 75.8 | 60.5 | -7.8 | 1.7 | 30 | 15.0 |
| R50_22 | 87.4 | 69.2 | -10.6 | 3.3 | 10 | 29.2 |
| R50_11 | 79.5 | 63.1 | -14.6 | -8. 7 | 0 | 6.2 |
| R50_01 | 83.7 | 66.2 | -15.9 | 14. 7 | 0 | 19.7 |
| R50_03 | 52.9 | 46.7 | -17.7 | -1 | 0 | 16.3 |
| R50_23 | 56.5 | 48.7 | -18. 6 | -17.3 | 45 | 15.3 |
| R50_09 | 57.1 | 49.1 | -26.9 | -16.3 | 30 | 8.9 |
| R50_18 | 73.6 | 59.1 | -27.4 | 22. 7 | 55 | 20.1 |
| R50_10 | 62.6 | 52.3 | -37.0 | -10 | 45 | 14.5 |
| R50_16* | 78.1 | 62.1 | 64.3 | 30 | 0 | 7.2 |
| R50_02* | 70.1 | 56.8 | -0.44 | 1.3 | 0 | 14.4 |
| Average | 76.5 | 62.0 | 5.8 | 12.1 | 24.9 | 13.35 |
| SD | 13.2 | 9.3 | 24.6 | 18.4 | 17.3 | 6.025 |
| SEM | 2.9 | 1.9 | 5.0 | 3.8 | 3.5 | 1.23 |
| Group: R30  Data provided for all 24 participants  Participants included in the analyses are indicated in grey (light grey across rows)  Positive values indicate reaching errors to the left of the target (in the expected direction of reach adaptation)  * outliers removed based on PEQ | | | | | |  |
| Pnum | Final Adaptation | Final Adaptation ARCSINE Transform | Process Dissociation Procedure (PDP) | Drawing Task (DT) | Post Experiment Questionnaire (PEQ) | Implicit (PDP) |
| R30_1 | 93.3 | 75.0 | 19.9 | 20.0 | 20.0 | 7.9 |
| R30_7 | 87.9 | 69.7 | 19.9 | 18.3 | 15.0 | 7.0 |
| R30_23 | 83.0 | 65.7 | 11.6 | 3.0 | 15.0 | 9.7 |
| R30_22 | 82.6 | 65.3 | 10.0 | -15.0 | 10.0 | 10.1 |
| R30_14 | 74.4 | 59.6 | 5.8 | -0.3 | 45.0 | 8.0 |
| R30_17 | 82.5 | 65.3 | 0.2 | 12.3 | 30.0 | 17.2 |
| R30_10 | 73.7 | 59.1 | -0.8 | -1.3 | 20.0 | 14.4 |
| R30_9 | 75.0 | 60.0 | -1.3 | -14.3 | 15.0 | 13.8 |
| R30_2 | 79.6 | 63.1 | -2.1 | 5.7 | 35.0 | 12.1 |
| R30_21 | 84.9 | 67.1 | -2.2 | -11.0 | 0.0 | 10.1 |
| R30_19 | 88.1 | 69.8 | -4.2 | 0.0 | 7.0 | 11.0 |
| R30_20 | 83.3 | 65.8 | -5.3 | -9.3 | 25.0 | 21.2 |
| R30_24 | 73.5 | 59.0 | - 8.3 | 4.7 | 45 | 16.3 |
| R30_15 | 87.7 | 69.5 | - 8.9 | - 0.3 | 15 | 14.0 |
| R30_06 | 86.1 | 68.1 | - 9.0 | 0 | 0 | 12.6 |
| R30_03 | 79.5 | 63.1 | - 9.5 | - 1.7 | 0 | 18.8 |
| R30_12 | 82.5 | 65.3 | - 10.0 | 3.7 | 20 | 12.9 |
| R30_18 | 80.4 | 63.8 | - 10.1 | 0 | 15 | 14.2 |
| R30_04 | 81.4 | 64.5 | - 10.8 | 0 | 10 | 10.6 |
| R30_05 | 77.4 | 61.6 | - 11.4 | 0 | 0 | 20.9 |
| R30_08 | 54.6 | 47.6 | - 16.5 | - 7.7 | 20 | 10.0 |
| R30_13 | 74.5 | 59.7 | - 29.1 | - 15 | 0 | 16.4 |
| R30_11* | 64.7 | 53.6 | 20.7 | -9.0 | 85.0 | 5.7 |
| R30_16* | 93.3 | 63.4 | 7.3 | 20.0 | 60.0 | 11.3 |
| Average | 79.6 | 63.5 | -1.84007962 | 0.72222222 | 21.125 | 12.7551 |
| SD | - 8.1 | 5.7 | 12.1644773 | 9.2316755 | 20.7349998 | 4.15 |
| SEM | -1.7 | 1.2 | 2.48306353 | 1.88440787 | 4.23251411 | 0.848 |
